# Restraint Stress Prolongs Diestrus Phase of Mouse Estrous Cycle

**DOI:** 10.1101/2025.11.07.687283

**Authors:** Gwendolyn R. Urbain, Alex D. Chapman, Brooke Van Loh, Joseph K. Folger, Geoffroy Laumet

## Abstract

Globally, stress levels among women of reproductive age are rising, while fertility rates continue to decline. Despite this correlation, a causal link between stress and reduced fertility remains unclear. Experimental studies have shown that severe and chronic stress can disrupt reproductive function, but the effects of mild stress, more representative of the daily stress experienced by most women, are still poorly understood. This study aims to identify how mild stress affects the mouse estrous cycle. Nineteen mice were vaginally lavaged daily one week before stress, during 3-day stress, and one week after stress. The mild stress paradigm consisted of two hours of repeated restraint stress each day for three days. Restraint stress disrupted the estrous cycle causing a longer cycle length in stressed mice, characterized by an extended duration in the diestrus phase. These findings suggest that even moderate stress perturbs normal reproductive cycling, potentially contributing to reduced fertility. This work highlights the need to further explore how everyday stressors may subtly impair reproductive health.

## Introduction

Fertility rates have been decreasing for the past decade (Datani, 2025). At the same time, women are reporting daily stress at an increasing rate (APA, 2023). Stress impacts women at a higher rate than men, a trend that might result from the dynamic neurochemical fluctuations in the female brain driven by cyclical hormonal changes (Lovick, 2012; Thiyagarajan et al., 2025). By using an animal model to study female cycling, external variables are controlled and eliminated such as biases/expectations about how stress impacts the body, patient, and psychological history (Caligioni, 2010). Rodents are commonly used to model the estrous cycle as their shorter reproductive cycle provides a quicker way to investigate changes, and estrous cycle tracking can serve as a proxy for the hormonal state of female mice (Rocks, 2022). Previously, chronic stress experiments have involved female mice being stressed over 30 consecutive days through cold water exposure, which caused decreased reproductive function (Casillas et al., 2021). Additionally, severe stress such as exposure to predator odors or 5-hours of restraint stress for 31 days caused an impaired luteinizing hormone (LH) surge, preventing ovulation from occurring (Wagenmaker et al., 2017). However, these extreme paradigms trigger life-threatening stress and may not appropriately model the daily stress experienced by average women. Therefore, there is an urgent need to address whether mild stress impacts the estrous cycle. In this project, a well-validated model of repeated restraint stress was used to mimic daily stress (Avona et al. 2020). We hypothesized that acute restraint stress would dysregulate the estrous cycle.

## Methods

### Animals

All animal procedures were approved by Michigan State University (MSU) Institutional Animal Care and Use Committee (IACUC) and in accordance with the National Institutes of Health (NIH) Guide for the Care and Use of Laboratory Animals. All animals were housed under a 12-hour light-dark cycle with *ad libitum* water and feeding. C57Bl/6J adult female (15 weeks old) mice were used as control (n=9) or stress (n=10) conditions. Mice were group housed (4-5 females per cage) for two weeks for acclimatization. The mice were randomly assigned to groups.

### Vaginal Lavaging

Mice were vaginally lavaged to collect vaginal epithelial cells by pipetting fifteen microliters of water onto the vaginal surface and removing the water. This process was repeated four times to ensure cell collection. The cells were then spread onto a slide in a circular fashion for even distribution.

### Staining and Staging

The slides with vaginal cells were stained using 0.5% Evans Blue dye and staged according to vaginal smear cytology rules (Cora et al. 2015). The smears consist of four phases: Proestrus (follicular phase), Estrus (ovulation), Metestrus (initial part of luteal phase), and Diestrus (later part of luteal phase) as illustrated in *Figure 1*. The mouse completed a full cycle by going through each phase in order.

**Figure 1:**
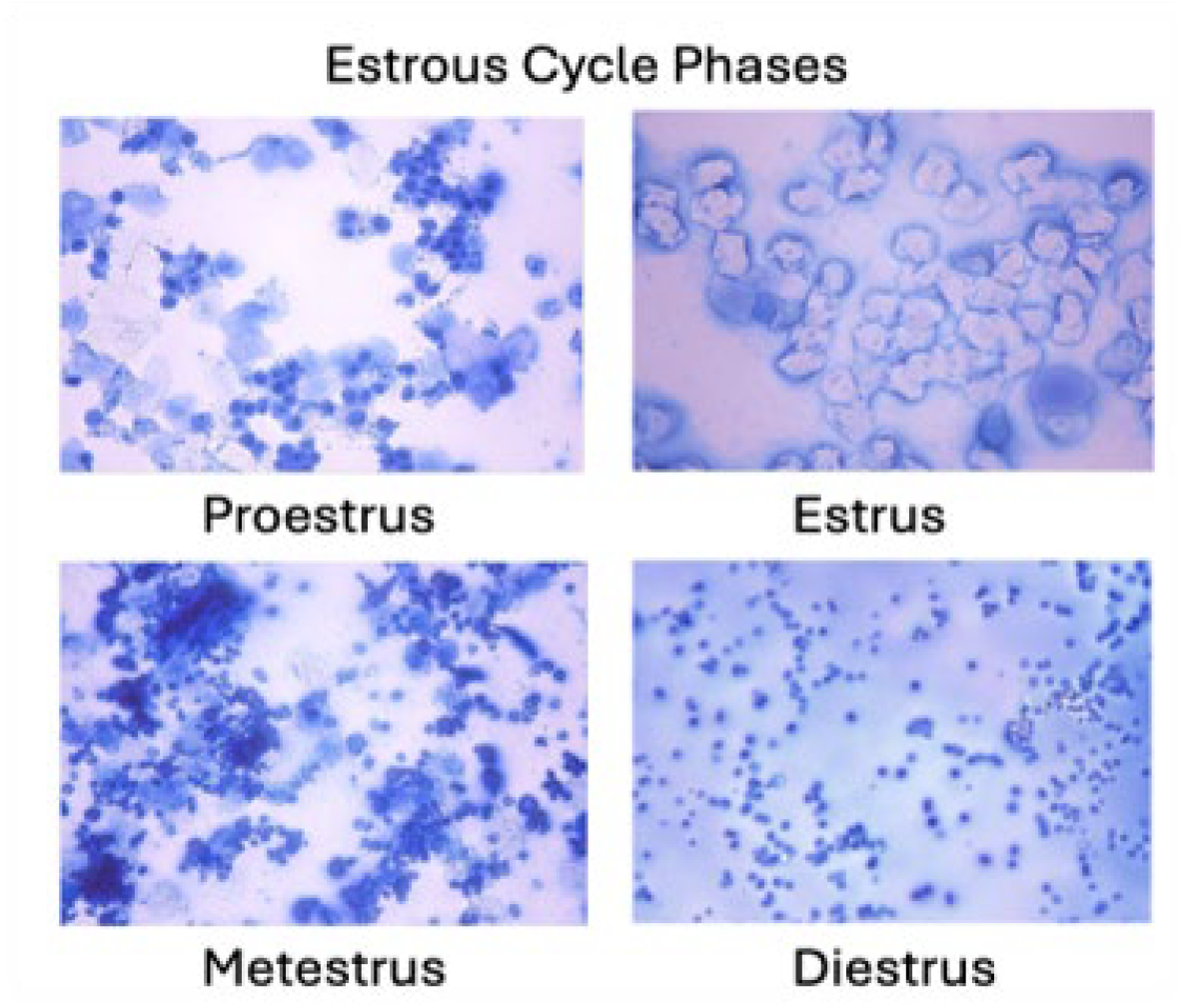
Estrous Cycle Phase determined by vaginal smears stained with 0.5% Evans Blue and staged using Brightfield Microscopy. **Proestrus**: Predominantly nucleated epithelial cells, indicating the onset of follicular development. **Estrus**: Abundant cornified (anucleate) epithelial cells, reflecting peak estrogen levels and sexual receptivity. **Metestrus**: Mixed cell population including leukocytes, nucleated, and cornified epithelial cells, marking the transition post-ovulation. **Diestrus**: Predominantly leukocytes, indicating low estradiol levels and luteal phase dominance.

### Restraint stress

Mice were placed into 50 ml conical tubes modified for restraint with breathing holes, and crumpled paper towels were inserted behind the mouse to hold it in place. Mice were restrained from 1100 h to 1300 h daily for 3 days.

### Timeline

The mice were lavaged daily at 1500 h for the entire 17-day period. This consisted of a pre-stress, during-stress, and post-stress timespan. The pre-stress phase consisted of seven days before stress to determine if the mice were normally cycling before stress. During stress, mice were restrained for two hours at 1100 h for three days. Control mice were left alone during stress period in home room. The post-stress period began at the end of stress and lasted for seven days. This timeline is represented in *Figure 2*.

**Figure 2:**
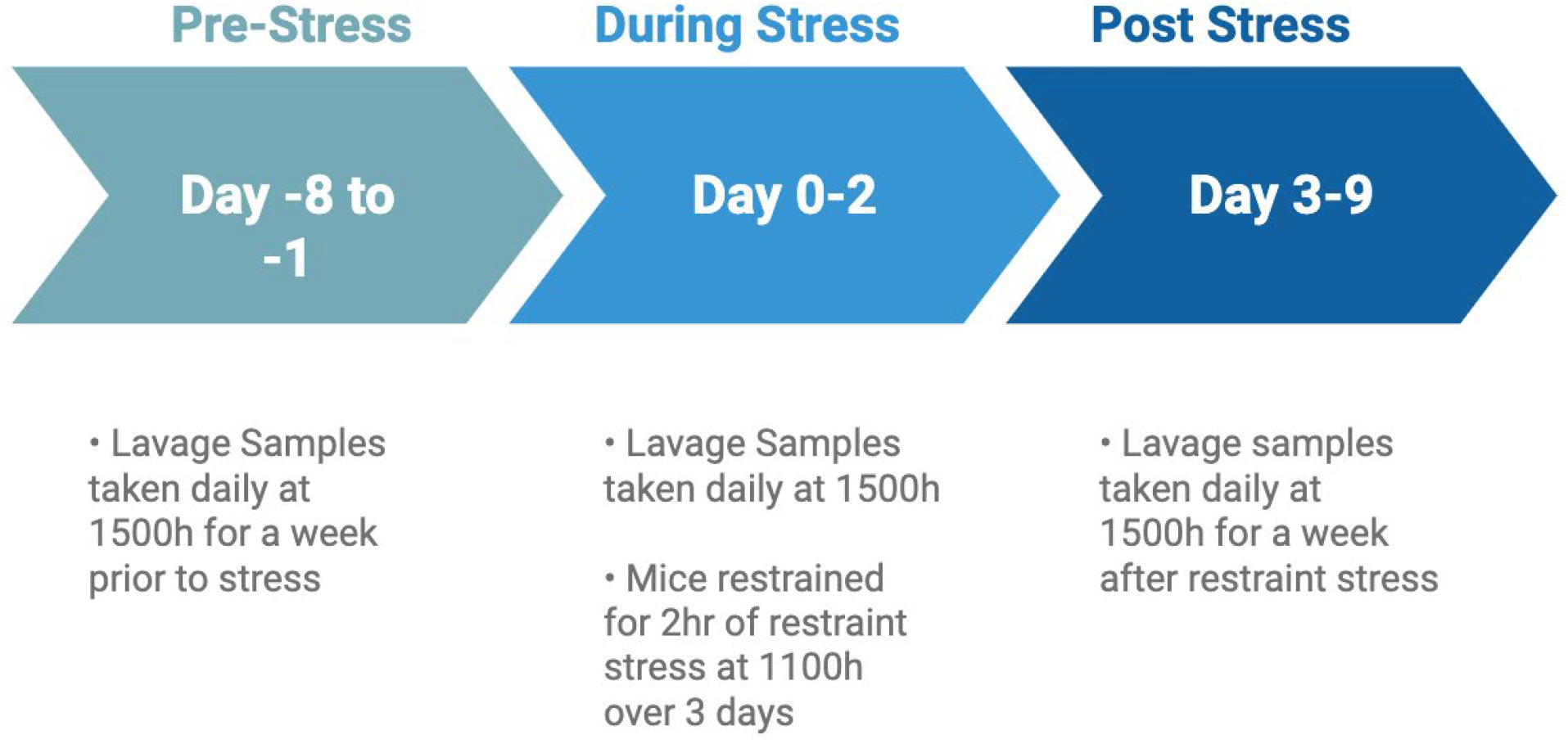
Experimental timeline of the 17 days consisting of pre-stress (Days −8 to −1), during stress (Days 0-2), and post-stress (Days 3-9) periods.

### Statistical Analysis

After staging slides for each mouse, each phase was assigned a number (Proestrus = 1, Estrus = 2, Metestrus = 3, Diestrus = 4) for quantification and graphical representation (*Table 1*). A mouse was counted as “cyclic” if it completed a full cycle (going through each phase in order i.e. 1→2→3→4, or 2→3→4→1, or 1→2→2→3→3→4 etc.) as seen by highlighted green cycles in *Table 1*. The number of cycles each mouse went through were then counted and put onto a separate table. Mice that did not complete a cycle during the post-stress period were assigned a cycle length of 10 days. The data was analyzed using GraphPad Prism v9. Based on experimental design, unpaired t-test, Fisher’s Exact test, or two-way ANOVA with correction for multiple comparisons were run to determine statistical difference between tested groups.

**Table 1:**
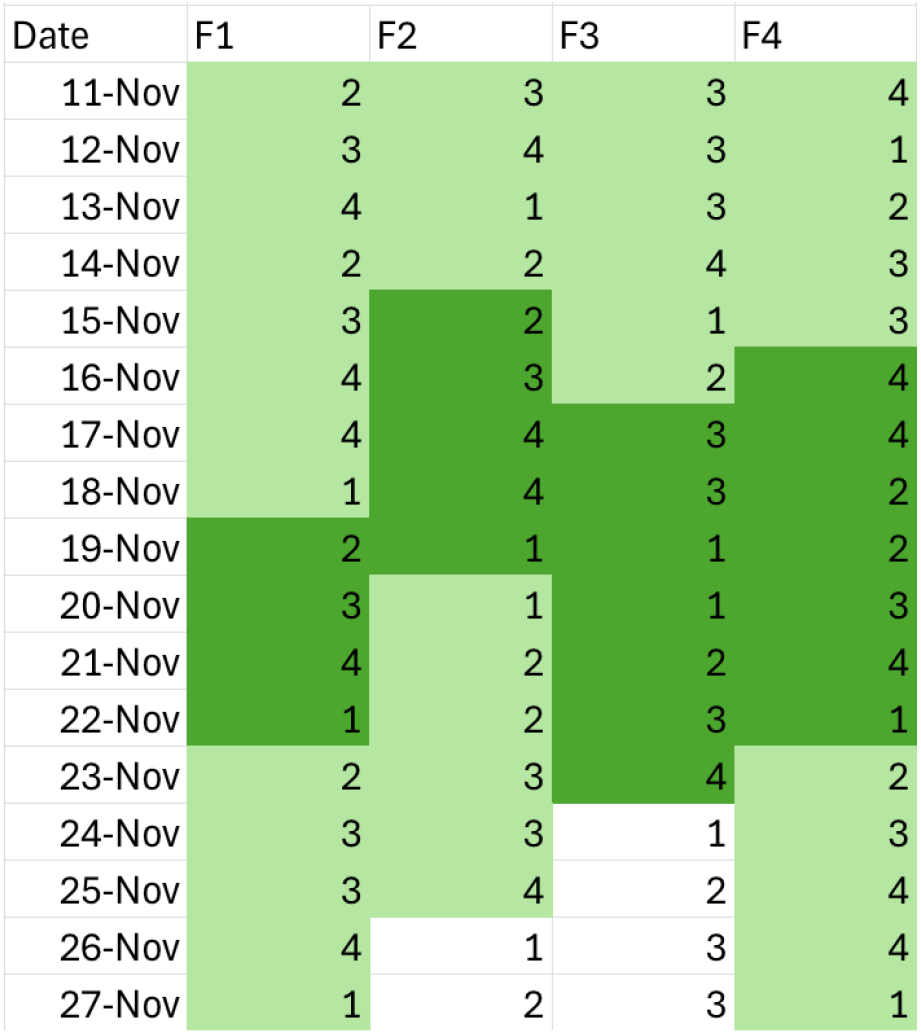
Representation of identified cycles in control mice. F1, F2, F3, and F4 represent females 1-4. For each day, the phase each mouse was in is identified and assigned a number (1-4 corresponding with estrous phase). A full cycle was highlighted in green. Different shades of green were used to differentiate cycles from one another.

| Date | F1 | F2 | F3 | F4 |
| --- | --- | --- | --- | --- |
| 11-Nov | 2 | 3 | 3 | 4 |
| 12-Nov | 3 | 4 | 3 | 1 |
| 13-Nov | 4 | 1 | 3 | 2 |
| 14-Nov | 2 | 2 | 4 | 3 |
| 15-Nov | 3 | 2 | 1 | 3 |
| 16-Nov | 4 | 3 | 2 | 4 |
| 17-Nov | 4 | 4 | 3 | 4 |
| 18-Nov | 1 | 4 | 3 | 2 |
| 19-Nov | 2 | 1 | 1 | 2 |
| 20-Nov | 3 | 1 | 1 | 3 |
| 21-Nov | 4 | 2 | 2 | 4 |
| 22-Nov | 1 | 2 | 3 | 1 |
| 23-Nov | 2 | 3 | 4 | 2 |
| 24-Nov | 3 | 3 | 1 | 3 |
| 25-Nov | 3 | 4 | 2 | 4 |
| 26-Nov | 4 | 1 | 3 | 4 |
| 27-Nov | 1 | 2 | 3 | 1 |

## Results

### Effect of Restraint Stress on Completion of the Estrous Cycle

All mice cycled normally for the 7-day pre-stress phase, before the start of our stress paradigm. As expected, in the control, 100% of female mice remained cyclic during the 10-day period of stress + post-stress. Restraint stress significantly reduced the number of completed cycles compared to control mice. (*Figure 3)*. In the stress condition only 40% of mice were cyclic after stress *(Figure 3)*. Given that fewer stressed mice completed a full cycle, potential impact of stress on the length of the cycle was examined. The control mice had an average cycle length of 5.9 days while stressed mice had an average cycle length of 8.8 days. The average length of the estrous cycle was significantly longer in the stress condition as compared to the control *(Figure 4)*. Additionally, 56% of the control group completed 2 full cycles, with the remaining 44% completing 1 full cycle over 10 days *(Figure 5)*. Only 40% of stress mice completed 1 full cycle, while 60% did not complete a full cycle over the 10-day period *(Figure 5)*.

**Figure 3:**
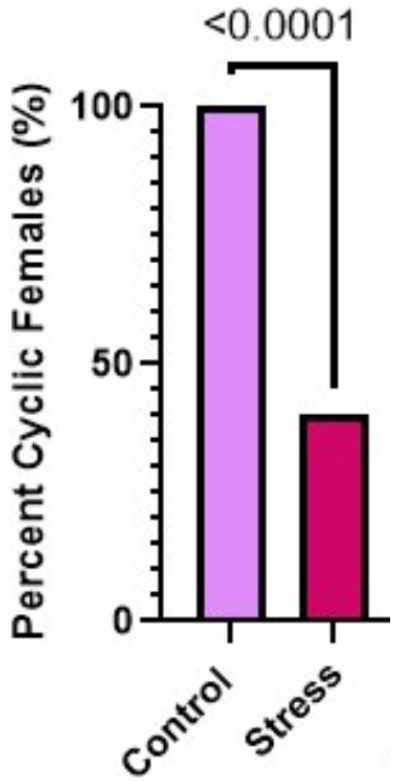
Restraint stress reduced the percentage of cyclic females (going through each phase in order). Mice were lavaged for 7 days after stress (n= 9 Ctrl and 11 stress). Fisher’s Exact test (**** p<0.0001)

**Figure 4:**
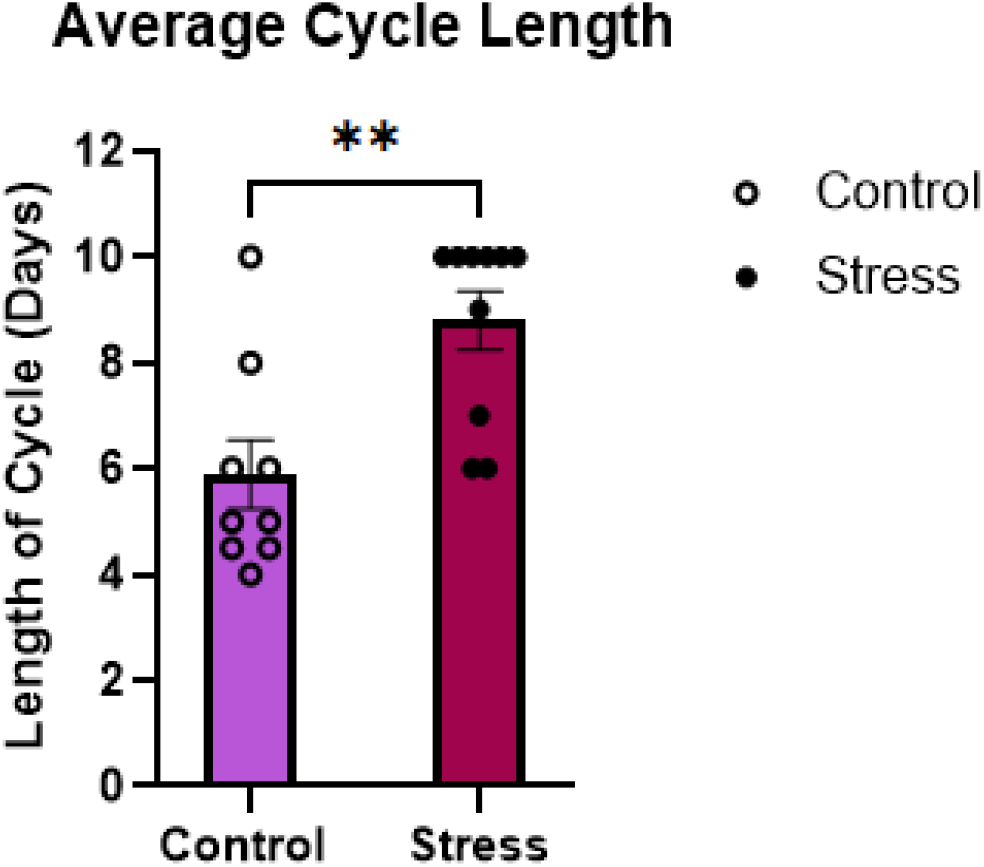
Restraint stress prolongs the length of Estrous Cycle. An unpaired T-test was used to compare control and stress conditions. Average cycle length was significantly increased in stress as compared to control. Mice that did not complete a cycle during the post-stress period were assigned a cycle length of 10 days. Unpaired T-test (F(8,9) = 1.24, **p<0.01).

**Figure 5:**
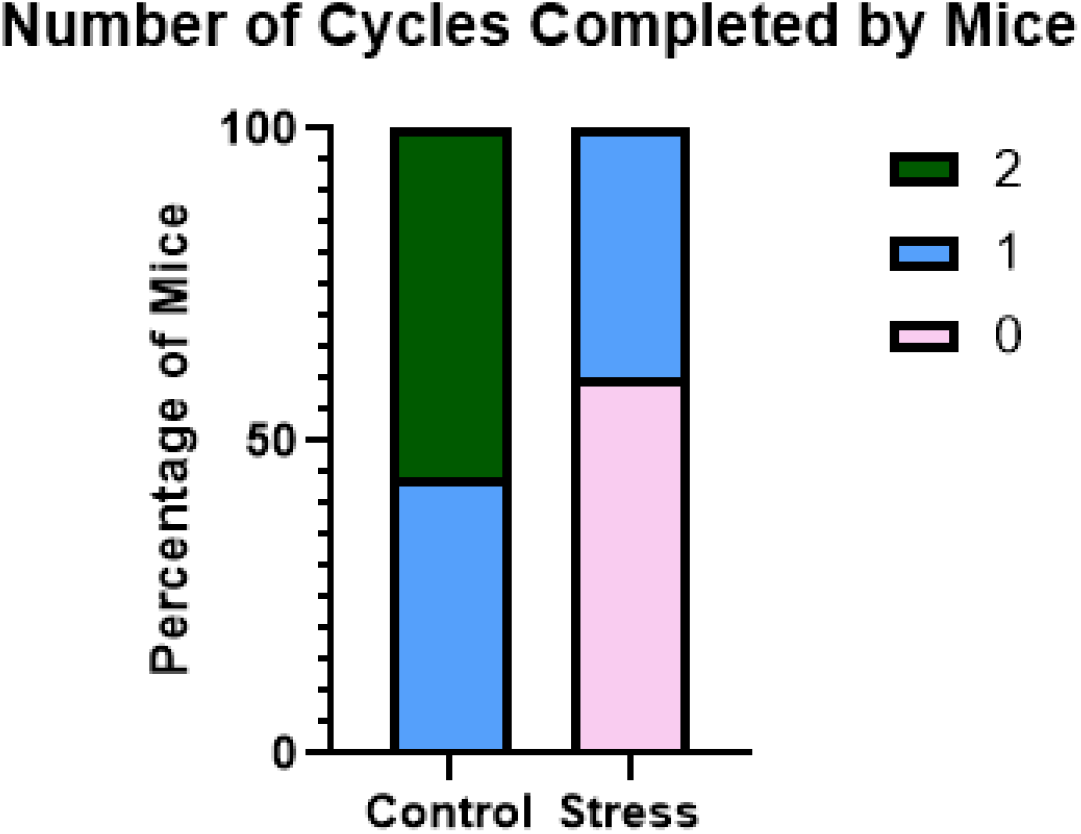
Restraint stress reduced the number of completed cycles over 10 days compared with stress and control conditions. Fisher’s Exact Test *** =p<0.001.

### Restraint Stress Causes Increased Time Spent in Diestrus Phase

As the stressed female mice completed fewer cycles and had longer cycle duration, the effect of stress on each phase was assessed. To determine whether a difference in phase length was present between stress and control mice, we measured the percentage of time spent in each phase. The pre-stress period showed all mice cycling normally with no difference in time spent in each phase between the control and stress mice *(Figure 6A)*. In control mice, the times spent in each phase stayed similar through the experiment *(Figure 6A, B, C)*. During the stress period, a significant decrease in time spent in the proestrus phase was seen in the stressed mice compared to control *(Figure 6B*). To assess whether stress was influencing a particular phase of the estrous cycle, we compared the time spent in each phase within groups. Stressed mice had a significant increase in time spent in the diestrus phase compared to all other phases during the stress period *(Figure 6B)*. After stress, mice significantly increased the time spent in the diestrus phase compared to control mice *(Figure 6C)*. Stress significantly elongated the diestrus phase, while other phases remained at the same length or shorter *(Figure 6C)*.

**Figure 6:**
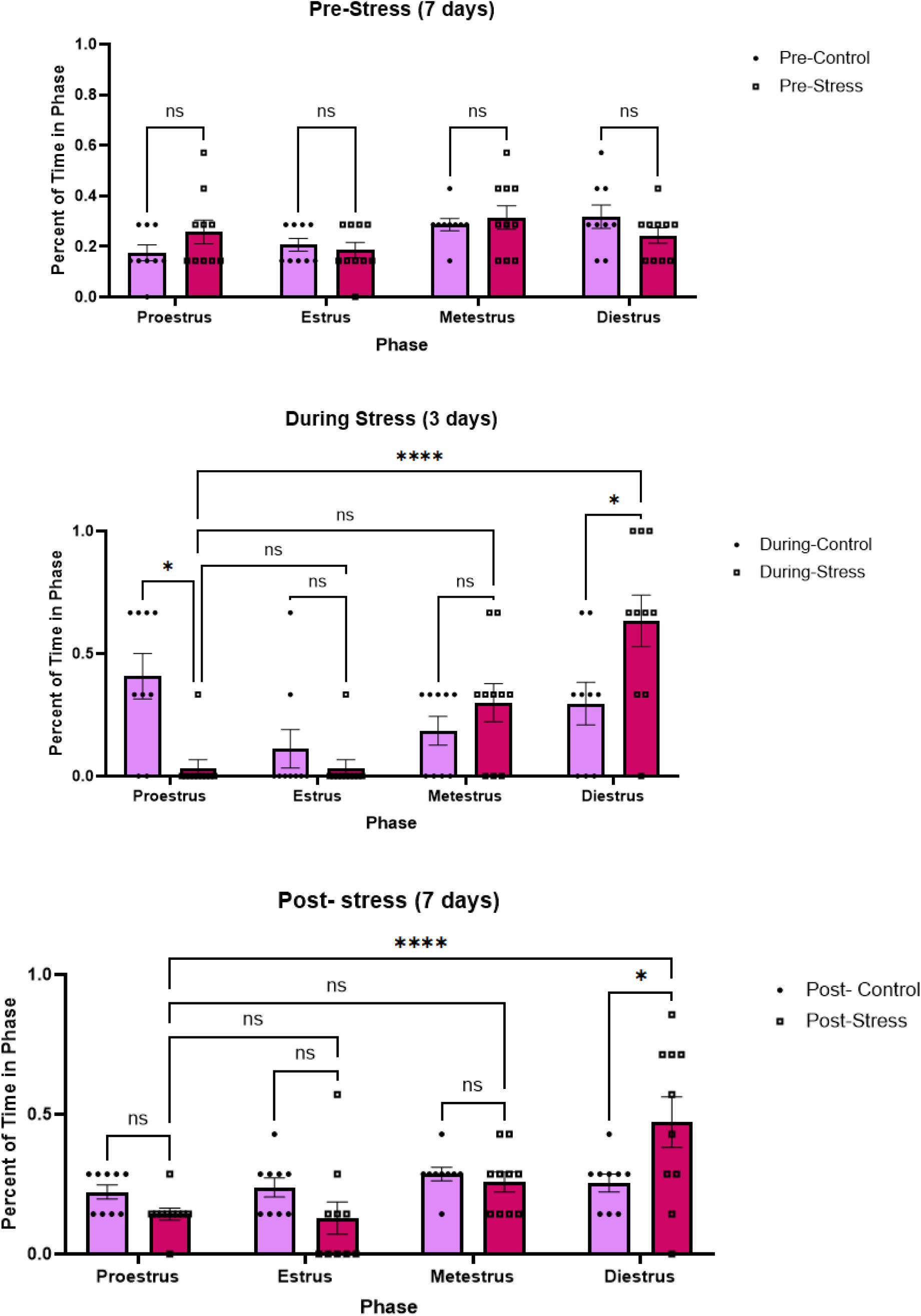
Restraint stress increased the percentage of time spent in diestrus phase and reduced the time in other estrous phases in mice. pre (A), during (B), and post (C) stress. Two-way ANOVA tests were used to determine the difference between stress and control conditions as well as within each condition’s phases. Two-way ANOVA: 6A: F(1,68) = 0.023; 6B: F(1,68) = 3.4e-19, F(3,68) = 9.43; 6C: F(1,68) = 2.01e-17, F(3,68) = 6.63, *p<0.05, **p<0.01, ***p<0.001, ****p<0.0001)

### Stress Mice Become Arrested in Diestrus Phase

To assess the effect of stress on estrous cyclicity, the temporal pattern of cycle phases was plotted to generate representative graphs for stressed and control mice. In the control condition, mice exhibited regular estrous cycles, progressing sequentially through each phase and completing two full cycles *(Figure 7)*. In contrast, mice exposed to stress showed disrupted cycling, completing zero to one full cycle *(Figure 7)*. Notably, stressed mice stayed in the diestrus phase, regardless of the phase they were in at the onset of stress *(Figure 7)*.

**Figure 7:**
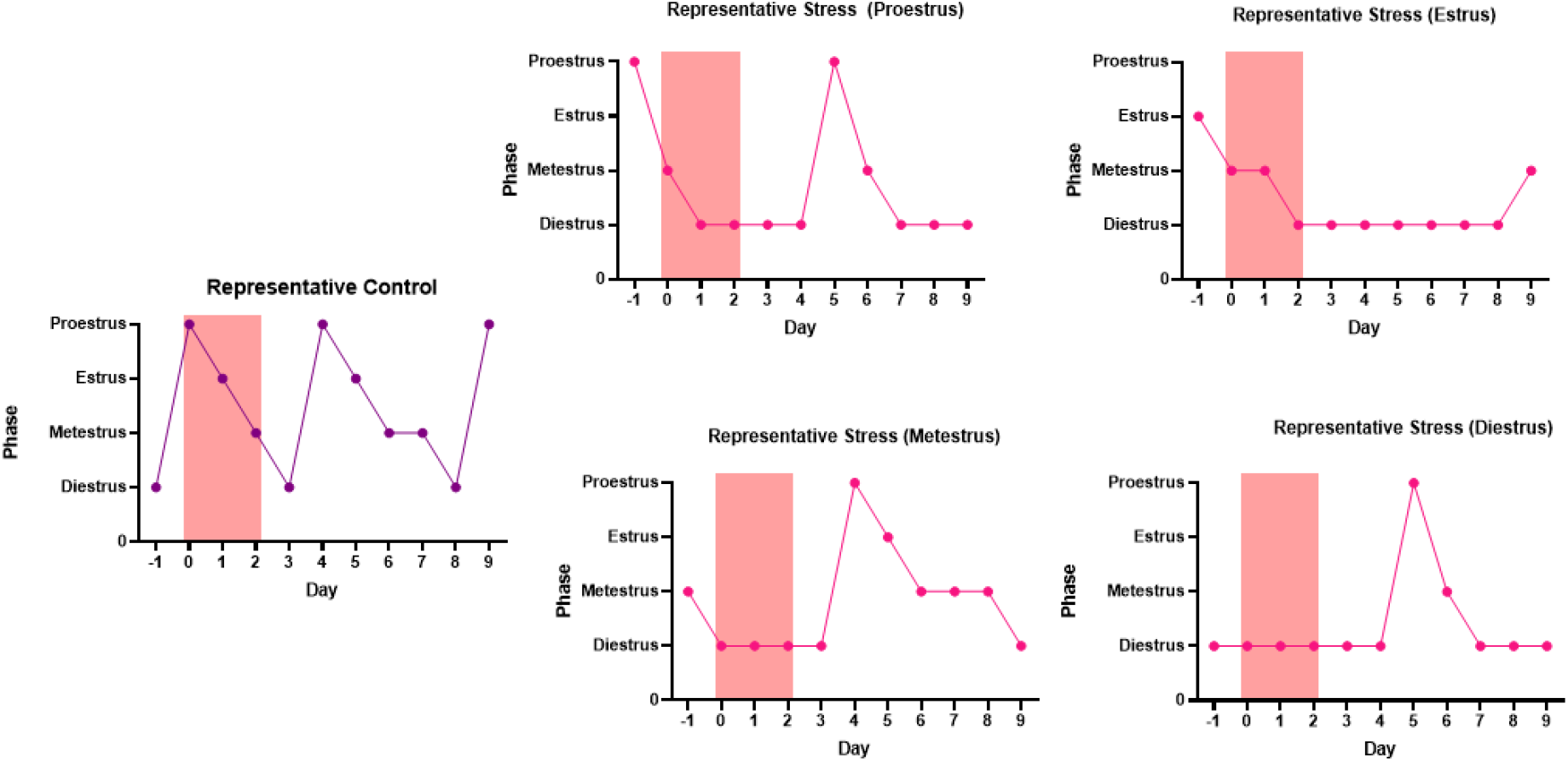
Representative graphs of daily phase tracking. Day 0 indicates the day the stress was initiated. Control can be seen in purple as the top graph. Stress period is represented by pink rectangle; control mice were not moved during this period.

## Discussion

This study examined how mild stress impacts the mouse estrous cycle using acute restraint stress. As fertility rates decline and women report stress at a higher level, identifying how stress impacts the female hormonal cycle becomes necessary (Datani, 2025; APA, 2023). This study showed a causal link between mild stress and compromised female estrous cyclicity. We determined that restraint stress alters the mouse estrous cycle by lengthening the cycle and increasing the amount of time spent in the diestrus phase. In all stressed mice, the diestrus phase was rapidly approached and markedly long. This confirmed our hypothesis that mild stress dysregulates the mouse estrous cycle and demonstrates a direct causal relationship between stress and female mouse reproductive health. Our findings reinforce the idea that stress stops cycling (Vigil et al. 2022).

Our findings show that acute restraint stress altered the mouse estrous cycle through lengthening of the estrous cycle. The elongation of the estrous cycle occurred regardless of the phase the mice began stress in, as seen in Figure 6. Because each mouse approached and was arrested in the diestrus phase, the data indicates that stress impaired estrous cycle regularity and specifically prolonged the diestrus phase. A lengthened diestrus phase could reinforce the idea that stress causes a state of lower neuronal signaling in the arcuate nucleus.

Studies have shown that acute psychosocial stress alters Kiss1 neuronal signaling which results in lowered Luteinizing Hormone (LH) levels due to increased RF-amide Related protein (Rfrp) gene expression (Yang et al. 2017). Rfrp acts as a Gonadotropin Inhibiting Hormone (GnIH) causing GnRH suppression. Inhibition of the pulsatile frequency of Gonadotropin Releasing Hormone (GnRH) leads to less production of Follicle Stimulating Hormone (FSH) and Luteinizing Hormone (LH) production (McCosh et al. 2023, Wagenmaker et al., 2008). Suppression of GnRH and LH pulses due to restraint (Yang et al. 2017) may explain that the cycle stalls in diestrus as a decrease in LH pulsation prevents the LH surge necessary for the cycle to continue. This means a full cycle is not observed. Without an LH surge, mice would be unable to reach ovulation, a key part of the reproductive process. By having dysregulated neuronal signaling, a female could be prevented from normal cycling, resulting in decreased fertility and impaired reproductive function.

Our understanding of stress and the female body is still limited, and studies focusing on hormonal interplay are greatly needed to understand potential fertility implications. Further research on how acute restraint stress impacts fertility aspects such as ovulation, pregnancy, pups per litter…, would increase our knowledge of the impact of stress on the female body. Animal models serve as an essential tool to determine the molecular mechanisms behind stress-related consequences to fertility.

## Acknowledgement and Conflict of Interest

This study was supported by the National Institute of Health (NIH) R01 NS121259, R01 AI177305, R21 NS142685 (G.L.), T32 GM142521 (A.D.C.), and the MSU College of Natural Science (G.R.U.). The authors declare no conflict of interest.

## Notes

### Competing Interest Statement

The authors have declared no competing interest.

## References

American Psychological Association. (2023, November). Stress in America 2023. American PsychologicalAssociation. https://www.apa.org/news/press/releases/stress/2023/collective-trauma-recovery

Avona A, Mason BN, Lackovic J, Wajahat N, Motina M, Quigley L, Burgos-Vega C, Moldovan Loomis C, Garcia-Martinez LF, Akopian AN, Price TJ, Dussor G. Repetitive stress in mice causes migraine-like behaviors and calcitonin gene-related peptide-dependent hyperalgesic priming to a migraine trigger. Pain. 2020 Nov;161(11):2539–2550. doi: 10.1097/j.pain.0000000000001953. PMID: 32541386; PMCID: PMC7572536.

Caligioni CS. Assessing reproductive status/stages in mice. Curr Protoc Neurosci. 2009 Jul; Appendix 4:Appendix 4I. doi: 10.1002/0471142301.nsa04is48. PMID: 19575469; PMCID: PMC2755182.

Casillas, F., Betancourt, M., Juárez-Rojas, L., Ducolomb, Y., López, A., Ávila-Quintero, A., Zamora, J., Ommati, M. M., & Retana-Márquez, S. (2021). Chronic Stress Detrimentally Affects In Vivo Maturation in Rat Oocytes and Oocyte Viability at All Phases of the Estrous Cycle. Animals, 11(9), 2478.

Cora MC, Kooistra L, Travlos G. Vaginal Cytology of the Laboratory Rat and Mouse: Review and Criteria for the Staging of the Estrous Cycle Using Stained Vaginal Smears. Toxicol Pathol. 2015 Aug;43(6):776–93. doi: 10.1177/0192623315570339. Epub 2015 Mar 3. PMID: 25739587; PMCID: PMC11504324.

Dattani Saloni, Rodés-Guirao L., and Roser M. (2025) - “Fertility Rate” Published online at OurWorldinData.org. Retrieved from: ‘https://ourworldindata.org/fertility-rate’ [Online Resource]

Driscoll AK, Hamilton BE. Effects of age specific fertility trends on overall fertility trends: United States, 1990–2023. National Vital Statistics Reports; vol 74 no 3. Hyattsville, MD: National Center for Health Statistics. 2025. DOI: 10.15620/cdc/174576.

Lovick TA. Estrous cycle and stress: influence of progesterone on the female brain. Braz J Med Biol Res. 2012 Apr;45(4):314–20. doi: 10.1590/s0100-879×2012007500044. Epub 2012 Mar 29. PMID: 22450372; PMCID: PMC3854171.

McCosh RB, O’Bryne KT, Karsch FJ, Breen KM. Regulation of the gonadotropin-releasing hormone neuron during stress. J Neuroendocrinol. 2022 May;34(5):e13098. doi: 10.1111/jne.13098. Epub 2022 Feb 6. PMID: 35128742; PMCID: PMC9232848.

Rocks D, Cham H, Kundakovic M. Why the estrous cycle matters for neuroscience. Biol Sex Differ. 2022 Oct 28;13(1):62. doi: 10.1186/s13293-022-00466-8. PMID: 36307876; PMCID: PMC9615204.

Thiyagarajan DK, Basit H, Jeanmonod R. Physiology, Menstrual Cycle. [Updated 2024 Sep 27]. In: StatPearls [Internet]. Treasure Island (FL): StatPearls Publishing; 2025 Jan-. Available from: https://www.ncbi.nlm.nih.gov/books/NBK500020/

Vigil P, Meléndez J, Soto H, Petkovic G, Bernal YA, Molina S. Chronic Stress and Ovulatory Dysfunction: Implications in Times of COVID-19. Front Glob Womens Health. 2022 May 23;3:866104. doi: 10.3389/fgwh.2022.866104. PMID: 35677754; PMCID: PMC9168655.

Wagenmaker ER, Breen KM, Oakley AE, Tilbrook AJ, Karsch FJ. Psychosocial stress inhibits amplitude of gonadotropin-releasing hormone pulses independent of cortisol action on the type II glucocorticoid receptor. Endocrinology. 2009 Feb;150(2):762–9. doi: 10.1210/en.2008-0757. Epub 2008 Oct 1. PMID: 18832098; PMCID: PMC2646534.

Wagenmaker ER, Moenter SM. Exposure to Acute Psychosocial Stress Disrupts the Luteinizing Hormone Surge Independent of Estrous Cycle Alterations in Female Mice. Endocrinology. 2017 Aug 1;158(8):2593–2602. doi: 10.1210/en.2017-00341. PMID: 28549157; PMCID: PMC5551545.

World Bank Open Data. Fertility rate, total (births per woman). (2023). https://data.worldbank.org/indicator/SP.DYN.TFRT.IN

Jennifer A Yang, Christopher I Song, Jessica K Hughes, Michael J Kreisman, Ruby A Parra, Daniel J Haisenleder, Alexander S Kauffman, Kellie M Breen, Acute Psychosocial Stress Inhibits LH Pulsatility and Kiss1 Neuronal Activation in Female Mice, Endocrinology, Volume 158, Issue 11, 1 November 2017, Pages 3716–3723, 10.1210/en.2017-00301

